# *Galleria mellonella* as an infection model for the multi-host pathogen *Streptococcus agalactiae* reflects hypervirulence of ST283

**DOI:** 10.1101/407171

**Authors:** Anne Six, Sakranmanee Kranjangwong, Margaret Crumlish, Ruth Zadoks, Daniel Walker

## Abstract

*Streptococcus agalactiae*, or group B streptococcus (GBS), infects diverse hosts including humans, economically important livestock and fishes. In the context of human health, GBS is a major cause of neonatal infections and an emerging cause of invasive disease in adults. Here we show that GBS is able to establish a systemic infection in *G. mellonella* larvae that is associated with extensive bacterial replication and dose dependent larval survival. This infection model is suitable for use with GBS isolates from both homeothermic and poikilothermic hosts and a hypervirulent sequence type (ST) associated with invasive human disease, ST283, shows increased virulence in this model, indicating it may be useful in studying GBS virulence determinants. In addition, we demonstrate that larval survival can be afforded by antibiotic treatment and so the model may also be useful in the development of novel anti-GBS strategies. The use of *G. mellonella* in GBS research has the potential to provide a low cost infection model that could reduce the number of vertebrates used in the study of GBS infection.

## Introduction

*Streptococcus agalactiae*, or Group B Streptococcus (GBS) is a Gram-positive multi-host pathogen and a major cause of human neonatal infections and mortality. GBS was initially described in 1887 as an animal pathogen responsible for mastitis in ruminants and although largely eradicated in some countries with a highly developed dairy industry, remains a major cause of bovine mastitis in many countries^1^. In 1966, GBS infection of fish was reported in the USA^2^ with subsequent outbreaks described in fish farms in Asia and the Americas^3,4^. GBS infection is often associated with high levels of mortality in fish and with the worldwide growth of aquaculture has significant economic impact^5^. In the 1970s GBS was described as the leading cause of human neonatal infectious morbidity and mortality with two distinct GBS-associated syndromes: early-onset disease occurring 0-6 days after birth and late-onset disease occurring 7 days - 3 months after birth^6,7^. The implementation of routine antibiotic prophylaxis during labour in many countries has resulted in a decrease of the incidence of early-onset disease, although the number of cases of late-onset disease has remained stable or even increased in some countries^7,8^.

GBS is also recognized as an emerging pathogen in adults that is associated with increasing fatality rates, reaching 15% in developed countries^9,10,11^. Adult GBS infections are often associated with underlying co-morbidities but a recent foodborne outbreak of invasive GBS disease in adults in Singapore was linked to the consumption of infected fish and occurred in previously healthy adults^12,13^. Despite some level of host adaptation, e.g. association of sequence type (ST) 67 with cattle^14^ and association of ST7, ST283 and clonal complex (CC) 552 with poikilothermic animals^15^, host-association is not absolute and interspecies transmission between people, cattle and fish may occur ^12,16,17^.

The routine use of antibiotic prophylaxis in pregnant women carrying GBS in many countries and routine use of antimicrobials in aquaculture and dairy farming may all exert selective pressure that favours emergence of antimicrobial resistance (AMR) in GBS^18^. Currently there is no GBS vaccine commercially available for human or bovine use and available fish vaccines do not confer protection against some serotypes. Consequently, the development of alternative anti-GBS strategies is of prime importance^19^.

A number of *in vivo* models exists for GBS infection (mice, rats, zebra fish) and are valuable for the study of host-pathogen interactions and virulence ^20,21,22^. In the last few years, larvae of the greater wax moth (*Galleria mellonella*) have been used as an alternate infection model for several human and non-human bacterial pathogens such as *Pseudomonas aeruginosa*^23^, *Bacillus cereus*^24^, *Listeria monocytogenes*^25,26^, *Vibrio anguillarum*^27^ and *Clostridium difficile*^28^. They have also been use to study the virulence of other *Streptococcus* spp.: *Streptococcus pyogenes*^29,30^, *Streptococcus pneumonia*^31^*, Streptococcus suis*^32^. Their ease of use, short life span and cost effectiveness make them a useful tool for screening of new antimicrobials and pathogen mutant libraries^33,34^.

Here, we evaluate the use of *Galleria mellonella* larvae as an *in vivo* model of infection for GBS by testing their susceptibility to human, bovine or fish GBS isolates encompassing host-associated and multi-host STs. We also determine the utility of *G. mellonella* larvae for *in vivo* screening of antimicrobial efficacy during GBS infection.

## Results

### G. mellonella larvae are susceptible to GBS infection

To begin to assess the suitability of *G. mellonella* larvae as an infection model for GBS strains isolated from its major hosts, we tested the susceptibility of larvae to dose-dependent killing by GBS strains from human, bovine and piscine origin. Larvae were challenged with serial dilutions of two isolates of ST17 (bovine strain MRI Z1-363, human MRI Z2-093) and one isolate of ST283 (piscine STIR CD25). Infected larvae were incubated at 37°C and survival was monitored 18h, 24h and 48h post-infection (**Figure 1A-C**). All three strains induced a dose-dependent response that was reproducible for each strain in three independent experiments. Immunohistochemical analysis of infected larvae with an anti-GBS antibody indicated extensive bacterial replication within larvae during the course of infection (**Figure 1D**). To generate quantitative data on the extent of GBS replication, CFU counts from homogenised infected larvae were measured for the piscine ST283 strain STIR CD25 (**Figure 1E**). CFU counts show extensive growth of ST283 up to 8 hours post infection, which is followed by a stationary phase of growth.

**Figure 1.**
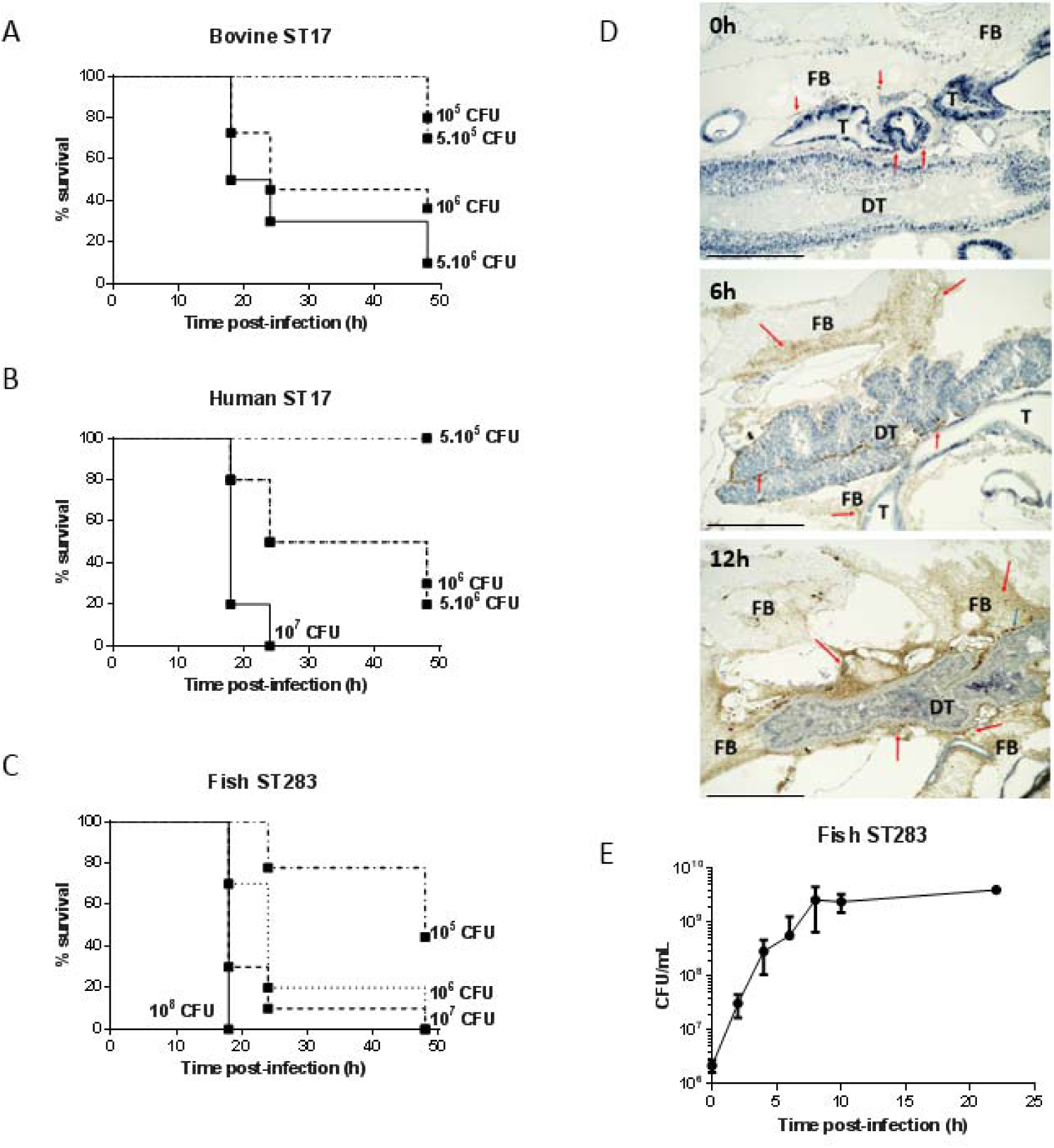
Virulence and replication of GBS in the *Galleria mellonella* larvae model. Kaplan-Meier survival curves of larvae challenged with serial dilutions of ST17 bovine isolate MRI Z1-363 (**A**), ST17 human isolate MRI Z2-093 (**B**), ST283 fish isolate STIR CD25 (**C**). **(D)** Immunohistochemical analysis shows extensive GBS replication in *G. melonella*. Larvae were inoculated with 10^6^ CFU of MRI Z2-366 (ST283) and collected immediately after inoculation and 6h and 12h post challenge. GBS were detected through immunohistochemical staining of sections. For histological examination, sections were stained with hematoxylin and eosin. Scale bar 500 µM. Abbreviations: DT, digestive tract; FB, fat body; T, trachea. Red arrows indicate the presence of GBS and blue arrows indicate phagocytic cells. (**E**) Bacterial counts of homogenized larvae showing *in vivo* growth of GBS ST283 isolate STIR CD25 in *G. mellonella* larvae following infection.

To further asses the utility of this model across a range of GBS strains, we tested the susceptibility of larvae to the fish specific ST260 isolate STIR CD10. Unlike piscine isolates of ST283, ST260 strains grow optimally at 28°C and does not grow at 37°C. Similar to experiments with non-ST260 strains at 37°C, STIR CD10 displayed dose dependent killing of *G. mellonella* larvae at 28°C (**Figure 2A**). To determine if incubation temperature is a critical parameter in determining the course of infection as has been previously described for a Galleria model of *Streptococcus pyogenes*^29^, one ST283 isolate (strain MRI Z2-366) was used to challenge larvae at both 28°C and 37°C. Percentage survival 24 h and 48 h post infection were similar at both temperatures, suggesting that the temperature of incubation post-challenge does not greatly influence virulence of GBS in this model (**Figure 2B**).

**Figure 2.**
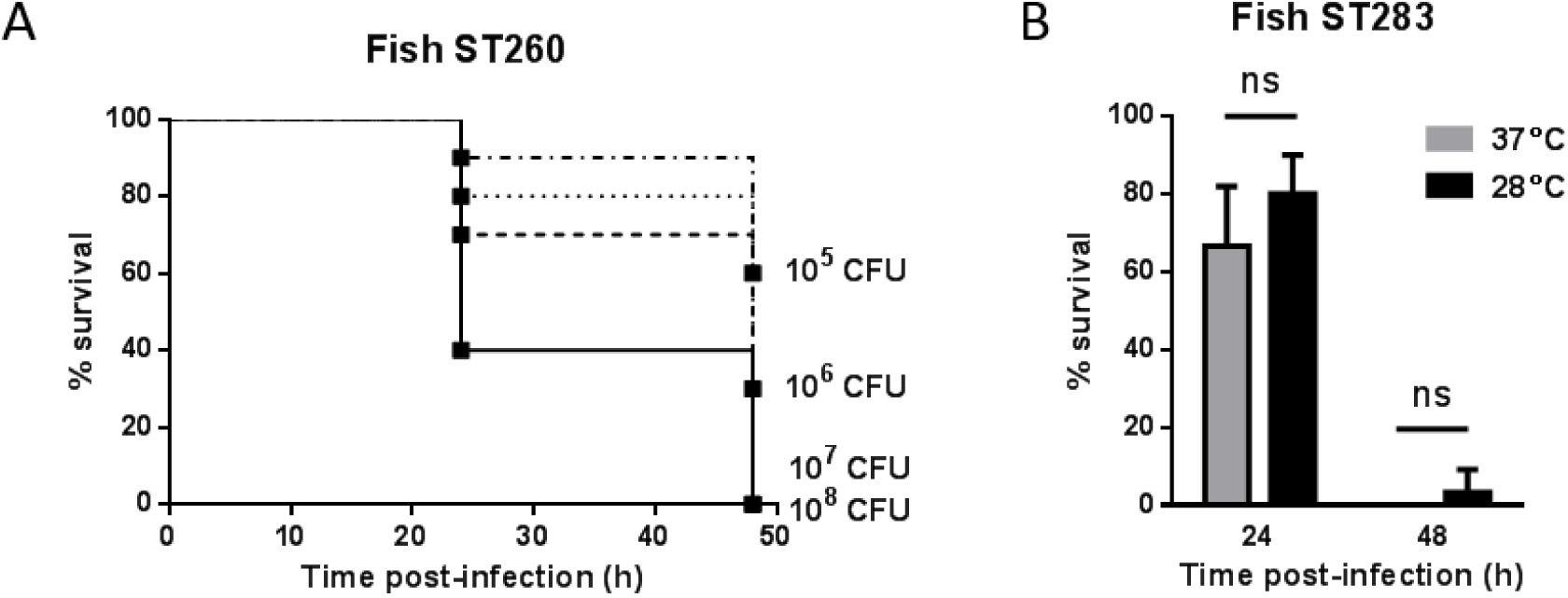
Virulence of GBS in the *G. mellonella* infection model is largely temperature independent. **(A)** Kaplan-Meier survival curves of larvae challenged with serial dilutions of ST-260 fish isolate STIR CD10 demonstrating dose dependent survival at 28°C. **(B)** Survival of larvae at 24h and 48h post-infection by ST283 fish isolate with incubation at 37°C or 28°C (ns : not significant; unpaired t-test).

### Invasive ST283 strains show increased virulence in G. melonella

Next, we examined if isolates from different host species showed different levels of virulence in *G. melonella* larvae through determination of the LD_50_ for GBS strains of human (n=9), bovine (n=12) and piscine origin (n=13). For each host species, we selected strains from the major epidemiologically relevant capsular serotypes and clonal complexes (**Table S1**). LD_50_ values for all isolates following infection of larvae at 37°C are shown in Table S1. Overall, LD_50_ values of human and bovine isolates were relatively homogenous with values ranging from 4.9 x 10^5^ to 3.2 x 10^6^ CFU and 1.1 x 10^6^ to 4.5 x 10^6^ CFU, respectively. Interestingly, LD_50_ values obtained for infection with fish isolates at 37 °C showed greater variance (ranging from 2.4 x 10^4^ to 1.1 x 10^7^ CFU) and were significantly lower than those of human and bovine isolates, indicating that fish isolates are more virulent in the *Galleria* model (**Figure 3A**). ST283 strains, which are responsible for a growing level of invasive GBS disease in healthy adults, generally showed the highest levels of virulence among GBS fish isolates. Comparison of LD_50_ values of ST283 and non-ST283 piscine isolates showed that there is a significant difference in virulence between these two groups (**Figure 3B**).

**Figure 3.**
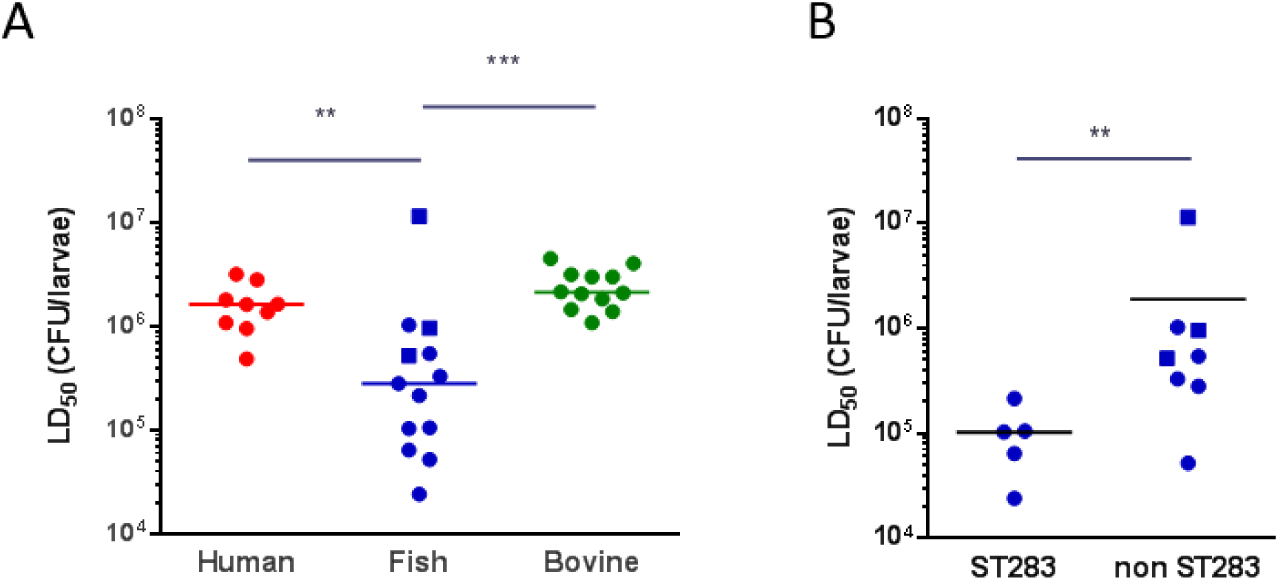
GBS isolates from fish, in particular ST283 isolates, show increased levels of virulence relative to human and bovine isolates. LD_50_ values determined by Probit analysis following infection of larvae by **(A)** *Streptococcus agalactiae* human (red), fish (blue) and bovine (green) isolates at 28°C (ST260 fish strains, square symbols) or 37°C (non ST260 strains, round symbols) and by **(B)** fish ST283 and non-ST283 isolates (** : p < 0.01; *** : p < 0.001; Mann-Whitney test).

### Antibiotic treatment rescues larvae infected with susceptible strains

To test the utility of the *Galleria* model to screen for antibiotic efficacy against GBS infection *in vivo* we tested the ability of erythromycin, ampicillin and tetracycline to afford survival against a lethal GBS infection. The three antibiotics were tested against 4 fish isolates (ST7, ST283, ST500, ST260), 2 human isolates (ST17, ST8) and 3 bovine isolates (ST8, ST7 and ST17). One ST260 fish isolate (STIR CD10) was tested by challenging the larvae and incubating at 28°C post-infection with the other strains tested at 37°C. Prior to infection studies, the MIC of each strain to erythromycin, ampicillin and tetracycline was determined by the broth dilution method (Table S1). All strains tested are susceptible to erythromycin and ampicillin with MICs ranging from 0.031 to 0.5 µg/ml. One fish (ST283; STIR CD25), one human (ST8; MRI Z2-137) and two bovine (ST8; MRI Z2-053 and ST7; MRI Z1-354) isolates were found to be resistant to tetracycline (MIC ranging from 16 to 64 µg/ml) while the other tested strains are susceptible (MICs ranging from 0.125 to 1 µg/ml). Except for those strains identified as resistant to tetracycline, one injection of 20 µg of erythromycin, ampicillin or tetracycline 2 hours post infection, was able to rescue larvae infected by fish, human or bovine isolates (**Figure 4**). For those strains resistant to tetracycline survival of Galleria was not significantly different on tetracycline treatment from control larvae treated with PBS. These data indicate an excellent correlation between GBS sensitivity to antibiotics and *in vitro* and *in vivo* at both 37°C and 28°C, indicating this model may be useful for testing novel anti-GBS agents.

**Figure 4.**
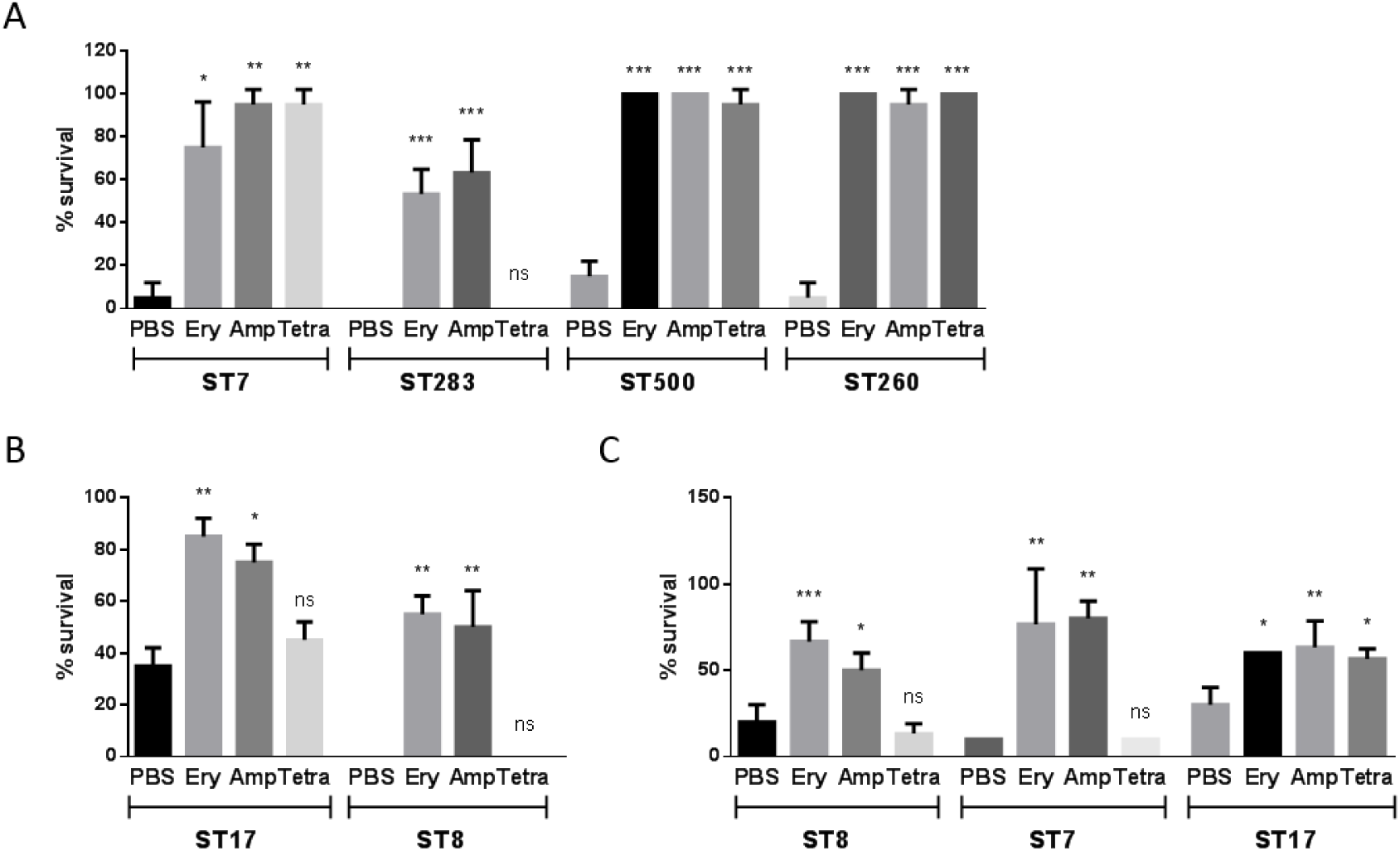
Antibiotic treatment affords survival of *G. mellonella* larvae against a lethal dose of GBS. Survival rates of treated (injection of 10 µL of 2 mg/mL erythromycin, ampicillin or tetracycline 2 hours post-infection) and untreated (10 µL of sterile PBS) larvae 48 hours post infection with GBS fish **(A)**, human **(B)** or bovine **(C)** isolates (ns: non significant, *: p < 0.05, ** : p < 0.01; *** : p < 0.001; one-way ANOVA).

## Discussion

In this work, we demonstrate the utility of *G. mellonella* larvae as a GBS infection model. GBS disease isolates are able to establish systemic infection of *G. mellonella* larvae with extensive bacterial replication and dose-dependent larval survival. The model is broadly applicable, with GBS isolates from multiple hosts, encompassing both homeothermic and poikilothermic animals, able to establish a lethal infection. Overall we observed that piscine isolates show increased virulence relative to human and bovine isolates. Interestingly, the most virulent fish isolates are largely strains of ST283, indicating that this hypervirulent ST may harbour virulence factors that contribute to infection in very diverse hosts including *G. mellonella*. This raises the possibility of utilising this cost effective model to screen transposon or other mutant libraries to identify key GBS virulence factors. However, some virulence factor (e.g. the lactose operon in bovine isolates) would not be reflected in a *Galleria* model and although correlation between the results obtained in the *G. mellonella* larvae model and other animal models have been shown for several pathogens^30,32^, disparity in the importance of virulence factors in different hosts has also been demonstrated^36^, highlighting the need for caution when searching for the impact of specific virulence factors in this model. Unlike other invertebrate models such as *Caenorhabditis elegans* and *Drosophila melanogaster*, *G. mellonella* larvae are able to survive at 37°C and 28°C, enabling the study of GBS infection in the range of relevant host-specific temperatures.

Due to the importance of GBS as a human, livestock and aquaculture pathogen and the widespread use of antibiotics to prevent or treat GBS infections there is a pressing need to identify and develop new anti-GBS strategies that could limit the associated spread of antimicrobial resistance. In this respect, invertebrate infection models are very attractive for the screening of new antimicrobials, to compare their efficacy to known antibiotics, or study the effect of antibiotic combinations^37,38,33,35^. To demonstrate the potential to use *G. mellonella* larvae for *in vivo* screening of antimicrobials we show antibiotics are able to afford survival to a lethal GBS infection. Indeed, the systemic injection of commonly used anti-GBS antibiotics, such as erythromycin, ampicillin or tetracycline, in the hemocoel of infected larvae is able to rescue those infected with susceptible strains but not those infected with resistant strains.

## Materials and methods

### Bacteria and growth conditions

GBS strains used in this study are described in Table 1. The panel of strains used represent a range of different capsular serotypes and sequence-types found in the three host species (fish, human and bovine). All human isolates originated from urinary tract infections so that potential differences in virulence between isolates within a host species could be associated with ST or serotype rather than clinical origin. Similarly, all bovine isolates originated from the milk of animals with mastitis. Where possible, human and bovine isolates of each ST were included to allow for assessment of host effects within ST. For fish isolates, the range of available STs was limited and represented the major CCs and serotypes known to be associated with fishes (ST7-serotype Ia; ST283-serotype III; CC552-serotype Ib). GBS were stored at -80°C in 25% (v/v) glycerol and cultured in Brain Heart Infusion broth/agar medium (BHI, Oxoid) at 37°C, with the exception of ST260, a member of the poikilothermic CC552, which were grown at 28°C.

### Galleria mellonella challenge

*G. mellonella* larvae were obtained from Livefood UK. They were kept in darkness at room temperature and were used maximum one week following arrival. Healthy larvae with no melanisation were used for all experiments. GBS strains were grown in BHI at 37°C or 28°C to an OD_600_ = 0.4 - 0.6. Cells were then washed twice in sterile PBS and diluted to the desired inoculum in PBS. Inoculums were serially diluted and plated on BHI agar plates just before administration for CFU counting. For the determination of the LD_50_ of GBS strains, 4 groups of 10 larvae were injected with 10 µL of serial dilutions of bacterial suspension in the hemocoel via the last right pro-limb. Following challenge, larvae were placed in an incubator at 37°C or at 28°C in the case of ST260 strains. Survival was followed for 48 hours, with health of surviving larvae assessed according to the health scoring index system [14]. Larvae were considered dead when non-responsive to touch. LD_50_ was calculated using the Probit method (XLSTAT software). Survival curves were plotted using the Kaplan-Meier method and differences in survival were calculated using the logrank test (GraphPad Prism 6). Experiments were done at least twice. One representative experiment was shown for survival curves and differences in LD_50_ between strains were assessed using Mann-Whitney test. To assess antibiotic efficiency, larvae were challenged with the LD_90_ of the GBS strain tested and injected 2 hours post infection with 10 µL of 2 mg/mL erythromycin, ampicillin or tetracycline in the hemocoel via the last left pro-limb. A sham-inoculation of PBS was used for untreated control larvae to account for mortality caused by physical injury or infection by a contaminant. Differences in survival rates between treated and untreated larvae were assessed using one-way ANOVA test followed by Dunnett test.

### In vivo GBS growth curve

For the *in vivo* growth curve experiment, larvae were challenged with 10^6^ CFU of GBS. At each time point groups of 3 larvae were kept at -20°C for 10 minutes before being transferred to Eppendorfs containing 100µL of sterile PBS, homogenized, serially diluted and plated on chromogenic selective GBS Brilliance agar (Thermo Scientific, PO5320A). Homogenization was performed using FastPrep lysing matrix M (MPbio).

### Immunohistochemistry (IHC) and histological analysis

Larvae were challenged with 10^6^ CFU of fish ST283 isolate (MRI Z2-366) as previously described. Uninfected and infected larvae were collected immediately after inoculation, 6h or 12h post challenge and kept at -20°C for up to 10 min before fixation. Whole larvae were placed in 10X volume of 10% neutral buffer formalin at room temperature for 48h. Longitudinal and cross sections of paraffin-embedded larvae were prepared to detect GBS presence in an early, middle and late phase of challenge. For antigen retrieval, heat induced epitope retrieval (HIER) was carried out using a Menarini Access Retrieval Unit (Biocare LLC, California, USA), in 10 mM Sodium Citrate buffer, pH 6.0 for 1 minute 40 seconds at 125°C full pressure. Slides were loaded on to Dako Autostainer (Dako Colorado, INC., Colorado, USA) and rinsed 5 min with a Tris-buffered saline solution (TBS), pH 7.6 containing 0.05% Tween 20. Endogenous peroxidase was blocked with Dako REAL(tm) Peroxidase-Blocking Solution (S2023) for 5 min and then rinsed with buffer. Slides were incubated 30 min at room temperature with anti-Streptococcus Group B antibody ab53584 (Abcam, Cambridge, UK) diluted 1: 200 in Dako universal diluent (S2022), washed and then further incubated with anti-rabbit horseradish-peroxidase labelled polymer (Dako, K5007ENV) for 30 min at room temperature. After incubation with 3,3’-diaminobenzidine (5 min, RT; Dako, K5007 DAB), the slides were rinsed three times with hydrogen peroxide and counterstained with Gills Haematoxylin for 27 seconds. Pictures were acquired on EVOS Fl Auto 2 Imaging System (Invitrogen).

### *Antibiotic*s

The minimum inhibitory concentration (MIC) was evaluated using the broth dilution technique for strains used in assessment of antibiotic efficiency. Microdilution plates containing 200 µL BHI broth medium with decreasing concentration of antibiotics (256 – 0.125 µg/mL) were inoculated from overnight cultures. After overnight incubation, cultures were checked for growth, with the MIC being the lowest concentration of antibiotics that prevents visible growth.

